# Genome-wide identification, characterization, and expression analysis of the monovalent cation-proton antiporter superfamily in maize, and functional analysis of its role in salt tolerance

**DOI:** 10.1101/2020.04.15.042580

**Authors:** Mengsi Kong, Meijie Luo, Jingna Li, Zhen Feng, Yunxia Zhang, Wei Song, Ruyang Zhang, Ronghuan Wang, Yuandong Wang, Jiuran Zhao, Yongsheng Tao, Yanxin Zhao

## Abstract

Na^+^, K^+^ and pH homeostasis are important for plant life and they are controlled by the monovalent cation proton antiporter (CPA) superfamily. The roles of *ZmCPAs* in salt tolerance are not fully elucidated. In this study, we identified 35 *ZmCPAs* comprising 13 Na^+^/H^+^ exchangers (*ZmNHXs*), 16 cation/H^+^ exchanger (*ZmCHXs*), and 6 K^+^ efflux antiporters (*ZmKEAs*). All ZmCPAs have transmembrane domains and maize protoplasts transient expression showed that ZmNHX5 and ZmCHX17 localized to cell membrane system. *ZmCHXs* were specifically highly expressed in anthers, while *ZmNHXs* and *ZmKEAs* showed high expression in various tissues. *ZmNHX5* and *ZmKEA2* were up-regulated in maize seedlings under both NaCl and KCl stresses. Yeast complementation experiments revealed the roles of *ZmNHX5, ZmKEA2* in NaCl tolerance. Analyses of the maize mutants further validated the salt tolerance functions of *ZmNHX5* and *ZmKEA2*. Our study highlights comprehensive information of *ZmCPAs* and provides new gene targets for salt tolerance maize breeding.

## 1. Introduction

High salinity is a significant abiotic stress factor restraining crop production worldwide. High concentration of sodium chloride (NaCl) or potassium chloride (KCl) affects the osmotic potential of the soil solution, thus inhibiting water absorption by plant. In addition, Na^+^ and Cl^-^ ions are easily absorbed to toxic levels by plant roots, which cause ion toxicity to plant cells [1, 2]. Under the above effects, salt stress eventually leads to plant metabolism disorder, oxidation-reduction system damage, photosynthesis limitation, and finally plant growth inhibition [3].

Many monovalent cation channels/transporters have been reported to play key roles in regulating salt tolerance in plants. In *Arabidopsis, AtNHX1*–6 and *AtNHX7*/*AtSOS1* were shown to be involved in maintaining Na^+^, K^+^ and pH homeostasis, and are critical for *Arabidopsis* salt tolerance [4, 5]. *AtKEA1, AtKEA3* and *AtKEA4* can enhance the tolerance of *Arabidopsis* to high concentration of K^+^, and *AtKEA2* overexpression enhanced the resistance of yeast mutants to Na^+^ and K^+^ stress. Functional characterization of *AtCHX17, AtCHX21* and *AtCHX23* knockout *Arabidopsis* mutants suggested their roles in K^+^ acquisition and in *Arabidopsis* salt tolerance. In rice, the expression of *OsNHX1*–*5* and *OsNHX7*/*OsSOS1* were increased by salt stress and they can suppress sensitivity of rice to high Na^+^ and K^+^ concentration [6, 7]. The K^+^/Rb^+^ competition assay of *OsCHX14* in *Xenopus laevis oocytes* verified its function in K^+^ homeostasis. In maize, the expression of *ZmNHX2* was up-regulated under NaCl stress in both shoot and root of maize seedling and it was presumed to be responsible for Na^+^ sequestration into vacuoles [8]; *ZmNHX7* putatively encodes a Na^+^ efflux plasma membrane protein and its expression increased in maize plants exposed to salt stress [9]. All these Na^+^/H^+^ exchangers (NHXs), K^+^ efflux antiporters (KEAs) and cation/H^+^ exchangers (CHXs) carried a conserved Na^+^/H^+^ exchanger protein domain (Pfam00999), and are members of the monovalent cation/proton antiporter (CPA) superfamily. However, the CPA gene family in maize has not been comprehensively studied, and the role of its members in salt tolerance have not been fully elucidated.

The CAP superfamily is grouped into the CPA1 and CPA2 families based on the Transporter Classification database (http://www.tcdb.org/) [10, 11] and the CPA1 family comprises the *NHXs* subfamily, while the CPA2 family is comprised of the *KEAs* and the *CHXs* subfamilies [4, 12]. The *NHXs* is predicted to contain 10–12 transmembrane (TM) domains. The *AtNHXs* consist of 8 members in *Arabidopsis*, and the *AtNHX1–4* and *AtNHX5–6* are localized inside cell, while *AtNHX7/AtSOS1* and *AtNHX8* are localized to the plasma membrane (PM) [4, 5]. The different subcellular location and tissue expression specificity between members are consistent with their different biological functions. For instance, the *AtNHX1–2* on vacuolar membrane are responsible for cell expansion in rapidly elongating organs; the *AtNHX5* and *AtNHX6* located at the trans-Golgi network are important for vesicle trafficking to the vacuole in different organs throughout plant development, while *AtNHX7* on PM affect membrane trafficking and vacuolar biogenesis in root cells [4, 5]. Compared with *Arabidopsis* and rice, the function of *ZmNHXs* in maize have not been well characterized.

The CPA2-type transporters have 8–14 putative TM domains [4]. The *Arabidopsis* had 6 members in the KEA subfamily [13]. The *AtKEA1* and *AtKEA2* are anchored in the inner envelope membrane of chloroplasts and are involved in chloroplasts division and membrane formation. The *AtKEA3* is located in the thylakoid membrane and it benefits photosynthesis efficiency [14, 15]. The *AtKEA4-6* were found mainly at the Golgi and they have a dramatic impact on K^+^, pH homeostasis in endomembrane compartments and in plant growth [4]. The function of *OsKEAs* in rice is rarely known. For the CHX subfamily, the *Arabidopsis* had 28 members. Among them, 18 *AtCHX* genes are pollen-specific or pollen-enhanced, and 6 are highly expressed in vegetative tissues, revealing the crucial role of *AtCHXs* in pollen development [4, 16]. In rice, *OsCHX14* was demonstrated to locate at Endoplasmic reticulum and regulate K^+^ homeostasis in flowering organs [17]. In maize, the members of *ZmKEAs* and *ZmCHXs* have not been investigated and their roles in salt tolerance need to be determined.

In this study, we conducted genome-wide identification of *ZmCPA* genes in maize. Their encoded proteins were classified into ZmNHX, ZmKEA and ZmCHX subfamilies, and their protein properties comprising molecular weight (MW), isoelectric point (pI), evolutionary relationships, number of TM domains, sub-cellular localization, and motifs were determined. The splice variants, exon–intron structure, and intron phases of identified genes were investigated. The transcriptional patterns of *ZmCPA* genes were analyzed during numerous developmental stages and under salinity stress. Six *ZmCPA* genes were functionally characterized by expression in a salt-sensitive yeast mutant. Analyses of two maize mutants, *zmnhx5* and *zmkea2*, further verified the functions of *ZmNHX5* and *ZmKEA2* in salt tolerance.

## 2. Results

### 2.1. Genome-wide identification and characterization of maize CPA superfamily genes

Extensive BLAST searches based on known *AtCPAs* identified a total of 35 ZmCPA superfamily proteins encoded in the genome of maize (Table S1). These were further confirmed through searching the Pfam database (http://pfam.xfam.org/search) for the existence of the signature Na^+^/H^+^ exchanger (PF00999) domain. The 35 genes were classified into three subfamilies: *ZmNHX* (13 genes), *ZmCHX* (16 genes) and *ZmKEA* (6 genes). The identified *CPA* genes were named *ZmNHX1–ZmNHX13, ZmCHX1–ZmCHX12, ZmCHX14–ZmCHX17, and ZmKEA1–ZmKEA6* according to previous reports or based on their degree of similarity with the family members in other species. The *ZmCPA*s were distributed on all chromosomes of maize, with 5, 3, 3, 4, 1, 3, 5, 4, 3, and 4 genes on chromosome 1, 2, 3, 4, 5, 6, 7, 8, 9, and 10, respectively (Fig. S1).

As most of the *ZmCPA* genes had more than one transcript annotated in the MaizeGDB database (Table S1 and Fig. S2). The transcripts with a conserved gene structure similar to that of their orthologs (*Arabidopsis*, rice, *Sorghum, Brachypodium distachyon*, and *Setaria italica*) were selected for further analyses. The transcript of each *ZmCPA* gene model was further validated using reverse transcription polymerase chain reaction (RT-PCR) assays with the gene-specific primers listed in Table S2. Notably, the transcripts of five *ZmCPA* genes verified by Sanger sequencing of cDNA were different from their annotation in MaizeGDB (Fig. S3).

The gene structures of the *ZmCPA* genes were determined by aligning their genomic sequences with cDNA sequences obtained by RT-PCR. In general, the *ZmCPA* genes in the same subfamily shared a conserved gene structure and the majority of introns in each family were in 0 phase (Fig. 1). The number of introns ranged from 11 to 25, from 0 to 5, and from 16 to 19 in *ZmNHXs, ZmCHXs* and *ZmKEAs*, respectively (Table 1). The average length of ZmNHX, ZmCHX, and ZmKEA proteins was 640, 824, and 815 amino acid (aa) residues, respectively. The average molecular weight (MW) of ZmNHX, ZmCHX, and ZmKEA proteins was 70.8, 88.4, and 87.4 kDa, respectively. The isoelectric point (pI) value of ZmNHX, ZmCHX, and ZmKEA proteins ranged from 5.28 to 9.07, from 5.95 to 9.88, and from 5.22 to 6.01, respectively (Table 1).

**Table 1.** Information of maize CPA genes identified in this study.

| Gene Name | Gene Model | Chromosomal Location | CDS (bp) | Protein Length (aa) |
| --- | --- | --- | --- | --- |
| *ZmNHX1* | *Zm00001d028330* | Chr1: 31041489-31049951 | 1599 | 532 |
| *ZmNHX7* | *Zm00001d031232* | Chr1: 182925112-182945175 | 3411 | 1136 |
| *ZmNHX11* | *Zm00001d003728* | Chr2: 56632216-56643296 | 2367 | 788 |
| *ZmNHX6* | *Zm00001d048732* | Chr4: 4469601-4475393 | 1641 | 546 |
| *ZmNHX8* | *Zm00001d019978* | Chr7: 81486383-81509317 | 1611 | 536 |
| *ZmNHX3* | *Zm00001d020892* | Chr7: 135679272-135687679 | 1491 | 496 |
| *ZmNHX10* | *Zm00001d021844* | Chr7: 164344371-164361391 | 2820 | 939 |
| *ZmNHX5* | *Zm00001d022504* | Chr7: 179047417-179052457 | 1620 | 539 |
| *ZmNHX4* | *Zm00001d045883* | Chr9: 45473833-45477106 | 1638 | 545 |
| *ZmNHX12* | *Zm00001d048459* | Chr9: 156705048-156709982 | 1440 | 479 |
| *ZmNHX2* | *Zm00001d024832* | Chr10: 90134129-90141743 | 1680 | 559 |
| *ZmNHX13* | *Zm00001d025052* | Chr10: 101802712-101807654 | 1563 | 520 |
| *ZmNHX9* | *Zm00001d026118* | Chr10: 138832710-138862665 | 2115 | 704 |
| *ZmCHX11* | *Zm00001d031077* | Chr1: 176192733-176196049 | 2388 | 795 |
| *ZmCHX15* | *Zm00001d031078* | Chr1: 176200721-176205899 | 2061 | 686 |
| *ZmCHX7* | *Zm00001d005032* | Chr2: 155039785-155042337 | 2553 | 850 |
| *ZmCHX4* | *Zm00001d041198* | Chr3: 105028259-105030854 | 2529 | 842 |
| *ZmCHX9* | *Zm00001d044623* | Chr3: 233503244-233505658 | 2415 | 804 |
| *ZmCHX8* | *Zm00001d049663* | Chr4: 39074035-39076527 | 2493 | 830 |
| *ZmCHX6* | *Zm00001d050509* | Chr4: 94925858-94928987 | 2232 | 743 |
| *ZmCHX2* | *Zm00001d053237* | Chr4: 221575337-221578465 | 2478 | 825 |
| *ZmCHX1* | *Zm00001d017805* | Chr5: 207302222-207308368 | 2484 | 827 |
| *ZmCHX12* | *Zm00001d035631* | Chr6: 37793506-37814621 | 2571 | 856 |
| *ZmCHX16* | *Zm00001d038517* | Chr6: 159098609-159101687 | 2598 | 865 |
| *ZmCHX3* | *Zm00001d021461* | Chr7: 152741608-152744549 | 2424 | 807 |
| *ZmCHX14* | *Zm00001d009889* | Chr8: 87464523-87467367 | 2391 | 796 |
| *ZmCHX5* | *Zm00001d010601* | Chr8: 121526743-121529490 | 2475 | 824 |
| *ZmCHX10* | *Zm00001d010629* | Chr8: 122278283-122281316 | 2874 | 957 |
| *ZmCHX17* | *Zm00001d012521* | Chr8: 176021482-176024817 | 2628 | 875 |
| *ZmKEA5* | *Zm00001d027466* | Chr1: 5893819-5906194 | 1791 | 596 |
| *ZmKEA4* | *Zm00001d001788* | Chr2: 935639-948444 | 3480 | 1159 |
| *ZmKEA3* | *Zm00001d041308* | Chr3: 111434917-111440097 | 2397 | 798 |
| *ZmKEA1* | *Zm00001d036981* | Chr6: 108202268-108211871 | 1848 | 615 |
| *ZmKEA6* | *Zm00001d046231* | Chr9: 73621310-73635421 | 1872 | 623 |
| *ZmKEA2* | *Zm00001d026645* | Chr10: 149326566-149340249 | 3315 | 1104 |

**Fig. 1.**
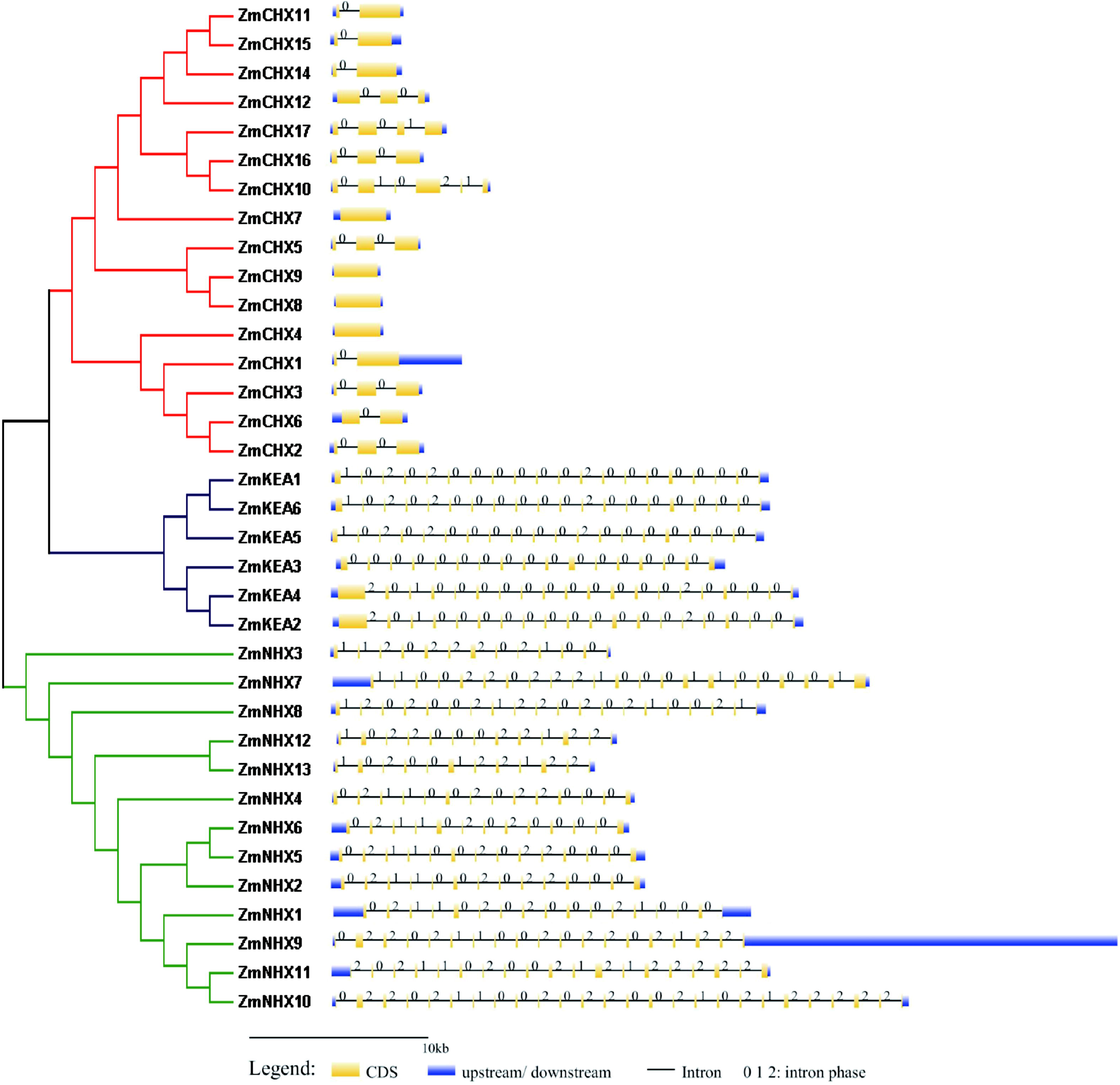
Gene structures of ZmCPA genes. Neighbor-joining tree of maize CPA genes at left is based on Fig. 3. Exon–intron organization of ZmCPA genes (right) as determined by aligning DNA and cDNA sequences of CPA genes at GSDS 2.0 server (http://gsds.gao-lab.org/index.php). Introns and exons are shown as thin lines and yellow boxes, respectively. Numbers 0, 1, and 2 represent different intron phases. Blue boxes at 5′ and 3′ ends represent untranslated regions (UTRs).

### 2.2. Phylogenetic analysis of maize CPA genes

To analyze the evolutionary relationships among the identified *ZmCPA* genes, we aligned their protein sequences with 26 OsCPA and 42 AtCPA proteins to construct a neighbor-joining (NJ) phylogenetic tree. Detailed information for *AtCPA* and *OsCPA* genes is shown in Table S3. Based on the topology of the phylogenetic tree, the *CPA* gene superfamily in plants can be subdivided into three subfamilies; NHX, CHX, and KEA. The NHX, CHX, KEA subfamilies can be further divided into groups N1*–*3, C1*–*3, and K1*–*2, respectively (Fig. 2). All groups contained genes from the three species of *Arabidopsis*, rice, and maize, indicating similar functions of orthologous genes in each group. We also performed a phylogenetic analysis using *ZmCPAs* only. As shown in Fig. 3, *ZmCPA* genes were grouped into *ZmNHX, ZmCHX*, and *ZmKEA* three subfamilies, which was consistent with the classification shown in Fig. 1 (Fig. 3).

**Fig. 2.**
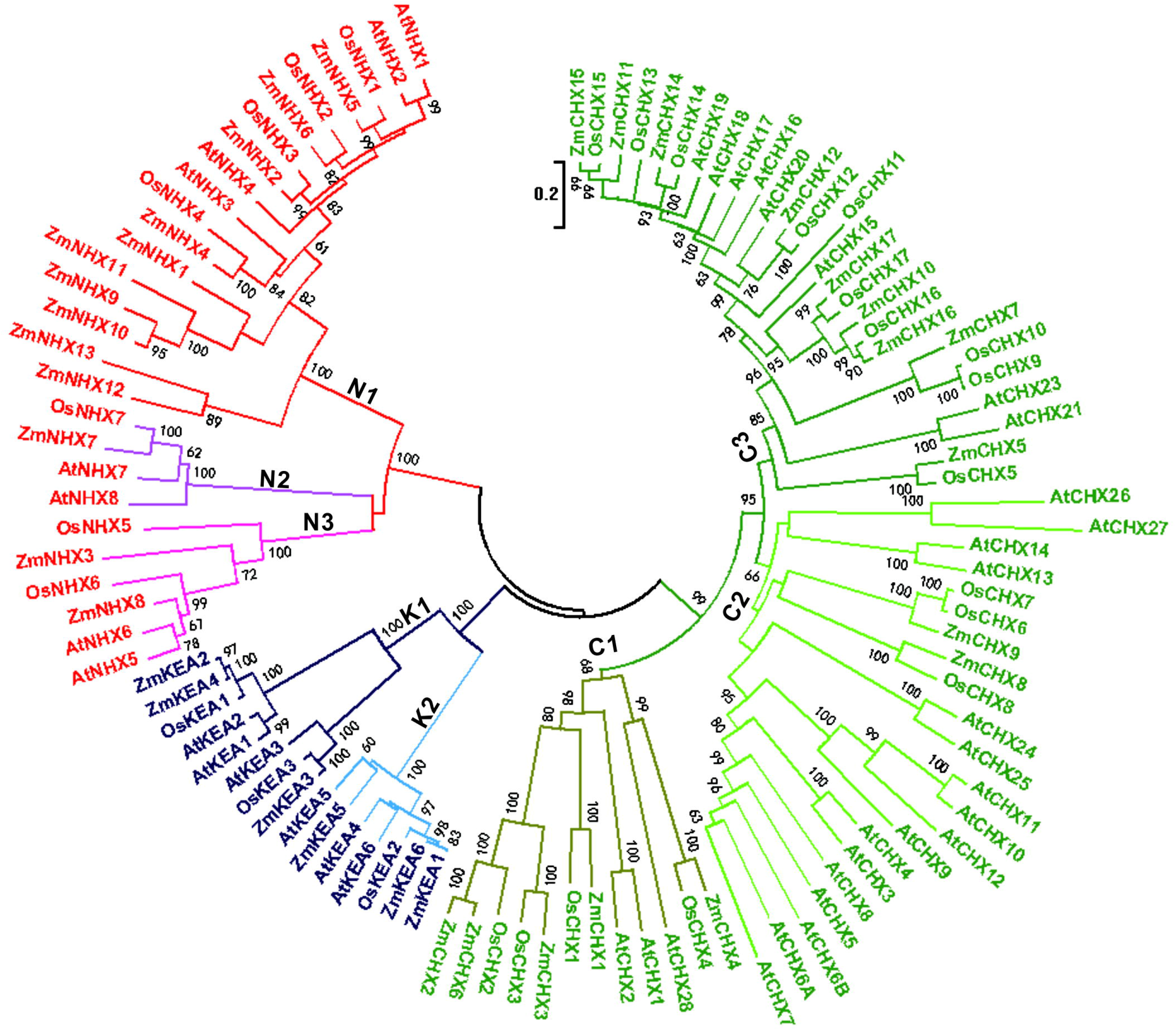
Phylogenetic tree of CPA genes in maize, Arabidopsis, and rice. Neighbor-joining tree constructed using MEGA 6.0 based on alignment of 103 CPA proteins including 35 ZmCPAs, 42 AtCPAs, and 26 OsCPAs. NHX, KEA, and CHX subfamilies are divided into three (N1, N2 and N3), three (C1, C2, and C3) and two (K1 and K2) groups, respectively.

**Fig. 3.**
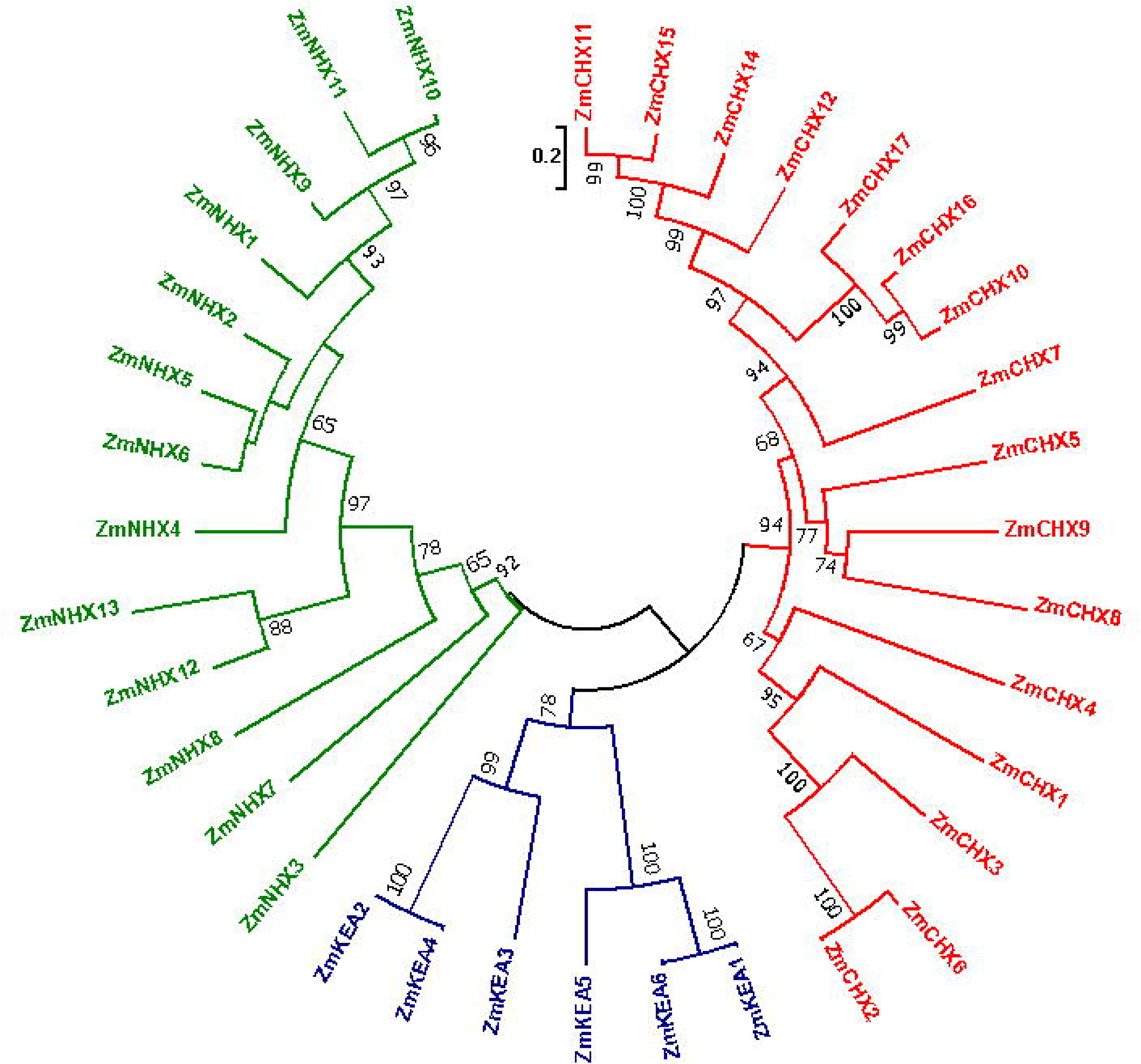
Phylogenetic tree of ZmCPA genes. All 35 ZmCPA proteins were aligned using MUSCLE v3.8.31 and alignment was used to construct neighbor-joining tree with MEGA 6.0. ZmCPA genes are divided into three subfamilies; ZmNHXs, ZmKEAs, and ZmCHXs.

### 2.3. Motif and subcellular localization analysis of ZmCPA proteins

Known conserved domains of ZmCPA proteins were identified by screening the Pfam database (http://pfam.xfam.org/). All proteins contained the Na^+^/H^+^ exchanger domain (PF00999), and three ZmKEAs and five ZmNHXs also contained the Trka_N domain (PF02254) and the TatD_DNase domain (PF01026), respectively (Fig. S4). Putative motifs of the ZmCPA proteins were mined using tools at the MEME server (http://meme-suite.org/tools/meme). The 15 motif sequences identified are listed in Table S4. Generally, ZmNHX proteins contained five to eight conserved motifs, the ZmCHX proteins had six to seven conserved motifs and the ZmKEA proteins had two to three conserved motifs (Fig. S4). Most closely related members in the same subfamily had common motifs, indicative of functional similarities among ZmCPA proteins.

The conserved domains of ZmCPA proteins corresponded to partial motifs and covered the TM domain (Figure S4*–*5). The TM helices within ZmCPA proteins were predicted using tools at the TMHMM Server v. 2.0 (http://www.cbs.dtu.dk/services/TMHMM/). All ZmCPA proteins had TM domains, and each subfamily had its own characteristics (Fig. S5). A maximum of 12 TM regions were predicted in ZmNHX5, -7, ZmCHX7, -16, -17, and ZmKEA1. The average occurrence of TM regions was 8, 10, and 10 in ZmNHX, ZmCHX and ZmKEA proteins, respectively (Table S1). The TM domain was mainly located at the C-terminal in ZmCHX subfamily, while at the N-terminal in ZmKEAs.

We performed transient expression assays in maize protoplasts to determine the subcellular localization of ZmCHX17 and ZmNHX5 proteins. The ZmNHX5-GFP and ZmCHX17-GFP fusion proteins co-localized with OsSCMP1-RFP protein (rice SECRETORY CARRIER MEMBRANE PROTEIN 1 fused with RFP as a membrane protein control [18]) suggesting that ZmNHX5 and ZmCHX17 localize to the membrane system (Fig. 4).

**Fig. 4.**
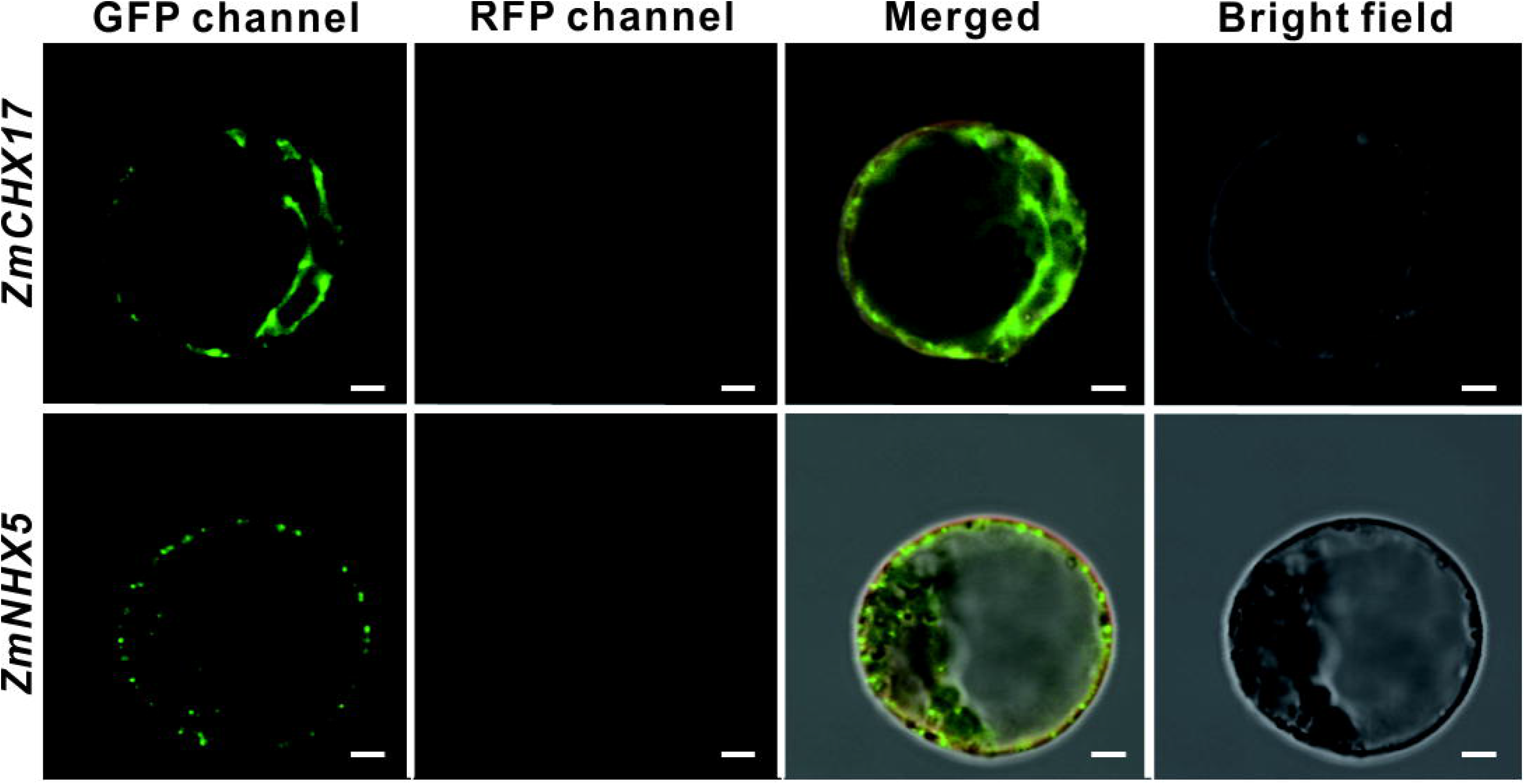
Subcellular localization of ZmCHX17 and ZmNHX5 proteins. ZmCHX17-GFP (upper panel) and ZmNHX5-GFP (lower panel) fusion proteins transiently expressed in maize protoplast cells co-transformed with OsSCAMP1-RFP fusion construct (membrane protein marker). GFP and RFP signals were detected by confocal fluorescent microscopy (Leica SP8). Scale bar = 5 μm.

### 2.4. Gene expression analysis in different tissues and developmental stages

NimbleGen maize microarray data [19] (ZM37) including 60 tissues representing 11 major organs and various developmental stages of the B73 maize inbred line were analyzed to detect the expression patterns of *ZmCPA* genes. Gene expression data for 31 *ZmCPA* genes including 10 *ZmNHXs*, 16 *ZmCHXs* and 5 *ZmKEAs* were used for the cluster analysis. As revealed by the heatmap, all *ZmNHX* genes except *ZmNHX4, ZmNHX3* and *ZmNHX9* were highly expressed in nearly all 60 tissues. All five *ZmKEA* genes were highly expressed in all 60 tissues. All *ZmCHX* genes were expressed at lower levels than *ZmNHX* and *ZmKEA* genes, and *ZmCHX* genes were only expressed at high levels in anther (Fig. 5).

**Fig. 5.**
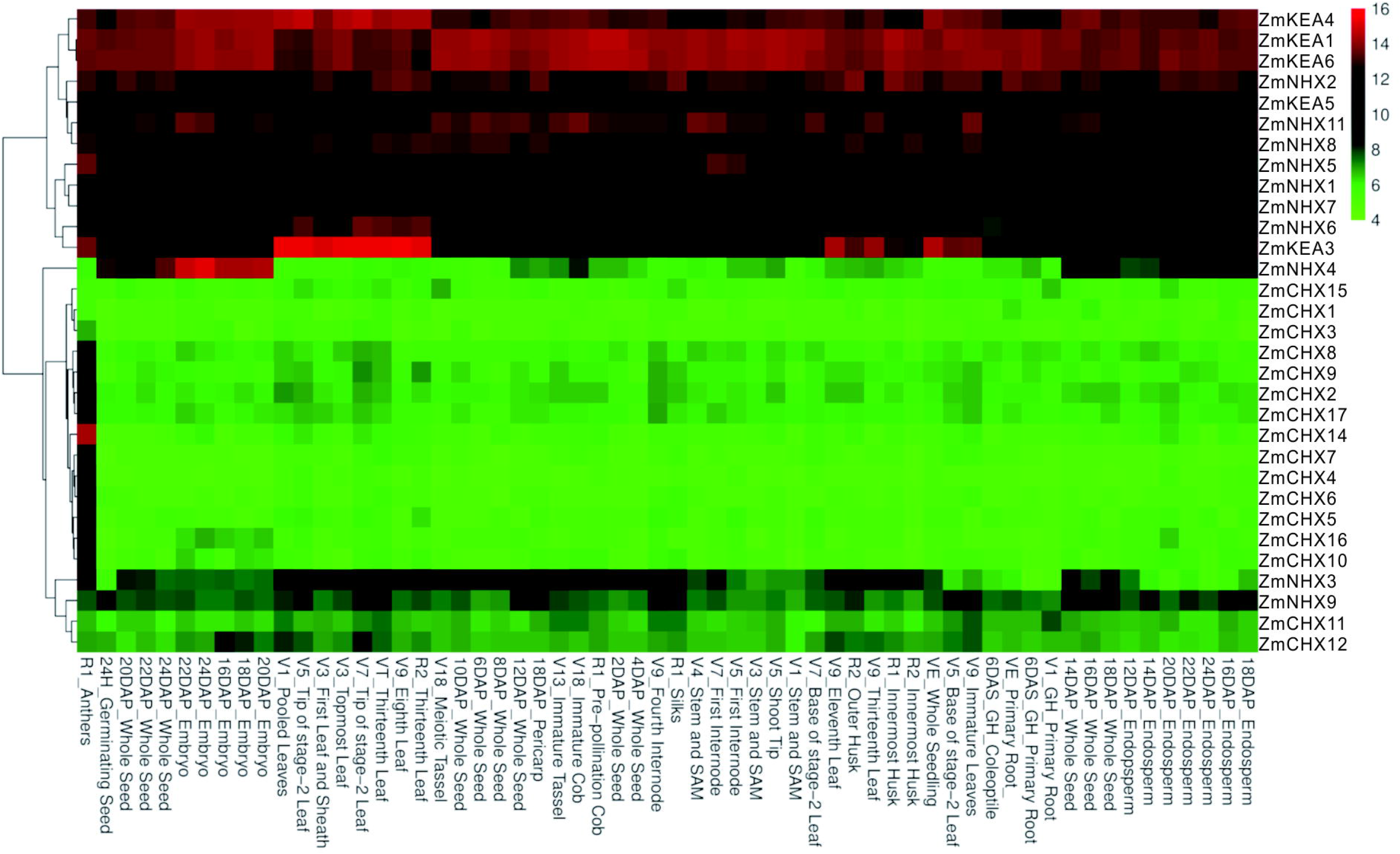
Gene expression profiles of ZmCPA genes in different tissues. Normalized microarray expression data of ZmCPA genes were downloaded from PLEXdb (http://www.plexdb.org/). Cluster analysis was performed and heatmap was generated using tools in the Lianchuan BioCloud Platform (https://www.lc-bio.cn/overview).

To confirm the tissue-specific expression of *ZmCPA* genes as revealed by the microarray data, semi-quantitative reverse transcription polymerase chain reaction (semi qRT-PCR) analyses of nine *ZmCPA* genes were performed with total RNA isolated from the roots, leaves, ears, immature tassels, pollen, anthers, silk, and whole seeds (20 days after pollination) of the B73 inbred line. The primers for semi qRT-PCR are listed in Table S5. Generally, the expression patterns of nine *ZmCPA* genes were consistent with the results from the microarray data. The *ZmNHX5, ZmNHX6* and *ZmNHX8* were highly expressed in root and leaf of maize, while the *ZmKEA1, ZmKEA2* and *ZmKEA5* were highly expressed in most of 8 tissues. The *ZmCHX8* showed low expression in all tissues, and the *ZmCHX5* together with *ZmCHX17* were specifically highly expressed in anther and pollen of maize (Fig. 5 and Fig. 6).

**Fig. 6.**
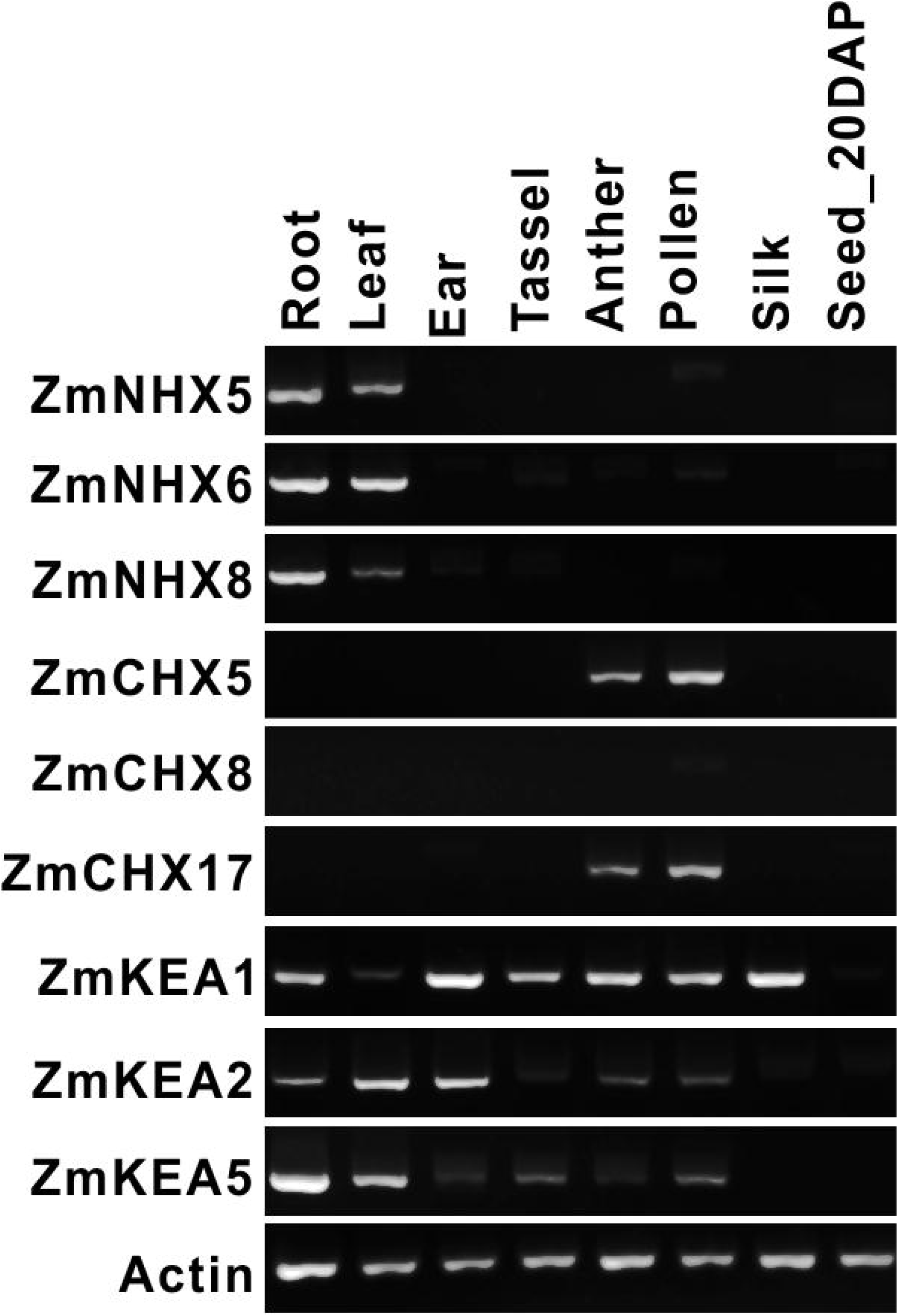
Expression analysis of ZmCPA genes in eight tissues of B73 inbred line by semi qRT-PCR. Total RNA was isolated from eight tissues: seedling roots, seedling leaves, 5-cm immature ears, tassels, anthers, pollen, silk and whole seeds (20 days after pollination, DAP), and then used for semi qRT-PCR analyses of ZmCPA genes. Maize Actin1 gene was the internal control.

### 2.5. Expression analysis of ZmCPA genes under 100 mM NaCl and 100 mM KCl stress

To understand the expressional responses of *ZmCPA* genes to salt stress, genes in two gene subfamilies, *ZmNHX* (four genes) and *ZmKEA* (four genes), were selected for analysis by real-time quantitative reverse transcription polymerase chain reaction (qRT-PCR) analyses. The primers are listed in Table S6. We detected the gene expression response at 0h, 1 h, 2 h, 4 h, and 24 h of salt stress. In the leaf of plants treated with 100 mM NaCl, *ZmNHX2, ZmNHX5, ZmNHX6, ZmKEA1* and *ZmKEA2* were up-regulated, while other genes were down-regulated. In the root, *ZmNHX5 and ZmKEA2* were up-regulated after 4 h, while other genes were down-regulated or did not change much. Under 100 mM KCl stress treatment, all all eight *ZmCPA*s (*ZmNHX2, ZmNHX5, ZmNHX6, ZmNHX8, ZmKEA1, ZmKEA2, ZmKEA5* and *ZmKEA6*) had a tendency of up-regulation in leaf. In the root, only *ZmNHX5* was up-regulated while other genes were down-regulated or showed fluctuation (Fig. 7). These results implied that *ZmCPA*s might play a role in salinity stress tolerance through expression regulation.

**Fig. 7.**
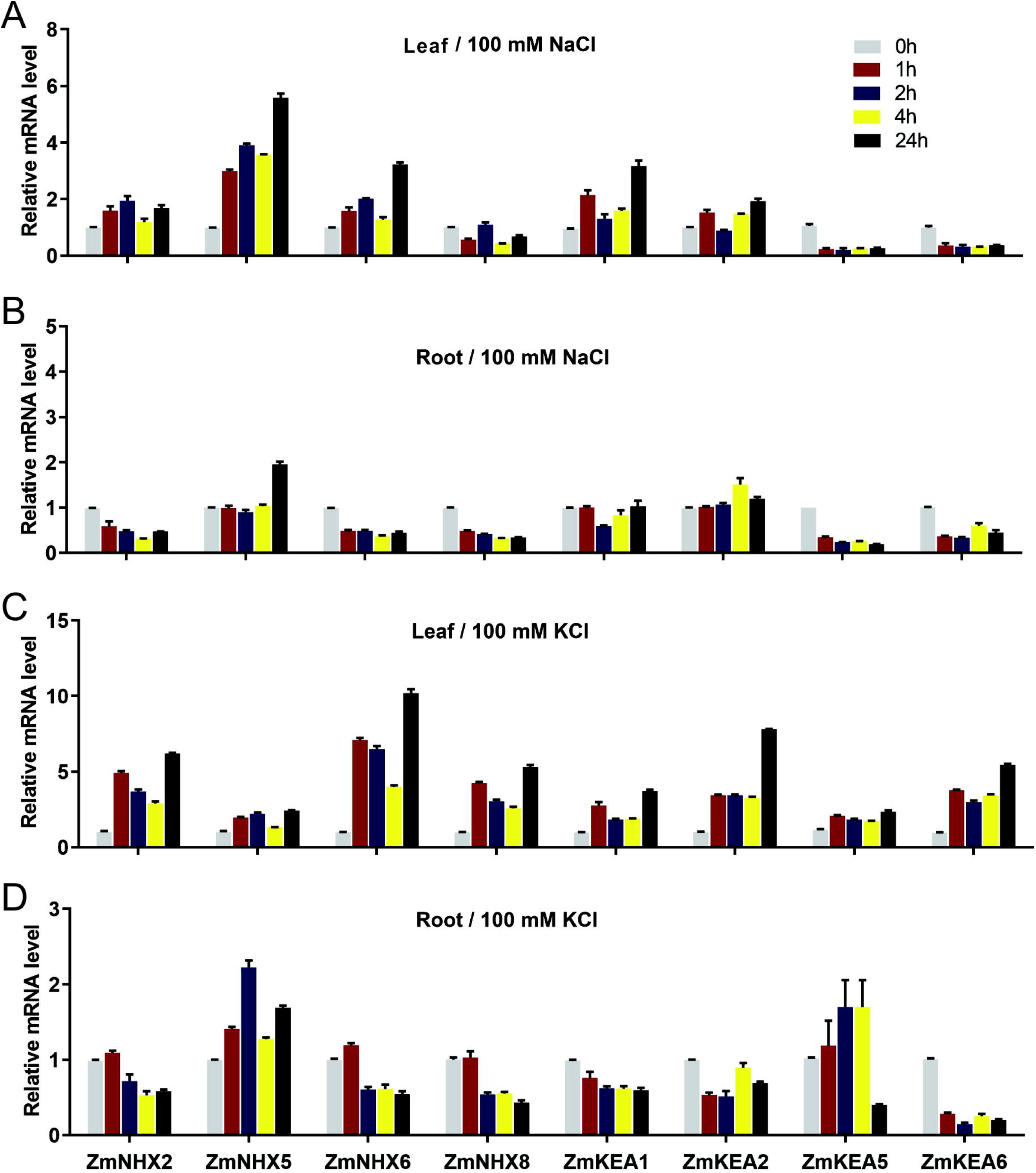
Expression analysis of ZmCPA genes under salt stress conditions by qRT-PCR. Expression analysis of ZmCPA genes under 100 mM NaCl or 100 mM KCl as determined by quantitative reverse transcription-polymerase chain reaction (qRT-PCR). Y-axis represents relative transcript levels of ZmCPA genes compared with that of actin1. X-axis represents different time points after salt treatment in each group. Error bars represent standard deviations for three replicates.

### 2.6. Functional analysis of ZmCPA genes in a yeast mutant under NaCl stress

To test the functions of *ZmCPA*s in salt tolerance, the coding sequences of *ZmNHX5, ZmCHX2, ZmCHX3, ZmCHX5*, and *ZmCHX17* and *ZmKEA2* were each cloned into the yeast expression vector pDR196 under the control of the PMA1 promoter. Each vector was introduced into the *Saccharomyces cerevisiae* mutant strain AXT3K, which lacks the functions of four plasma membrane Na^+^-ATPases (ScENA1–4), the plasma membrane Na^+^, K^+^/H^+^ antiporter ScNHA1, as well as the vacuolar Na^+^, K^+^/H^+^ antiporter ScNHX1 [20]. Therefore, this mutant is very sensitive to Na^+^ stress. Each transformed yeast strain was grown on Arg phosphate (AP) medium with or without 20 mM NaCl. Results showed that the transformed AXT3K lines expressing these genes grew much better than the AXT3K mutant as well as the AXT3K mutant transformed with the empty pDR196 vector under 20 mM NaCl stress (Fig. 8). These results confirmed the functions of *ZmNHX5, ZmCHX2, ZmCHX3, ZmCHX5*, and *ZmCHX17* and *ZmKEA2* in salt tolerance.

**Fig. 8.**
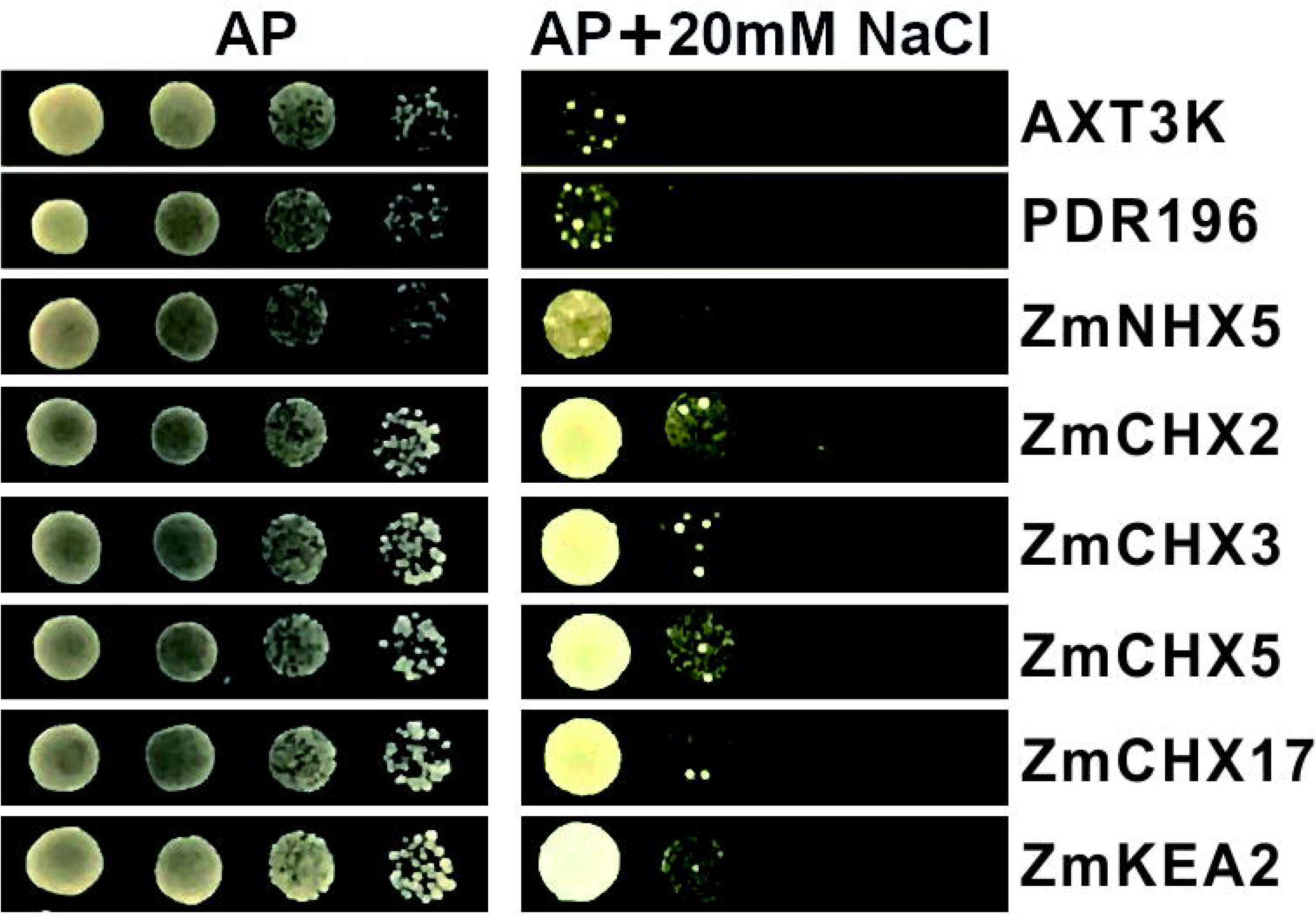
ZmCPA genes facilitate growth of yeast mutant strain AXT3K under salt stress. Full-length CDS sequences of ZmNHX5, ZmCHX2, ZmCHX3, ZmCHX5, ZmCHX17, ZmKEA2 were each cloned into PDR196 vector, then transformed into yeast strain AXT3K (ena1-4::HIS3,nha1::LEU2, nhx1::KanMX). Yeast transformant cells were cultured overnight, then were diluted with water to A_600_ of 0.8. Aliquots (5 μL) of 10-fold serial dilutions were spotted on AP plates supplemented with 20 mM NaCl at pH 7.5 (right panel) and normal AP plates as the control (left panel). Strains were grown at 29°C for 5 days and photographed.

### 2.7. Functional analysis of ZmNHX5 and ZmKEA2 under high concentration salt stress

We obtained two maize mutants, *zmnhx5* (EMS4-0a18d8) *and zmkea2* (EMS4-02c2af), which were produced by EMS mutagenesis of the B73 inbred line, from the Maize EMS-induced Mutant Database (MEMD) [21]. The *zmnhx5* and *zmkea2* mutants have pre-termination mutations in *ZmNHX5* (*Zm00001d022504*) and *ZmKEA2* (*Zm00001d026645*), respectively, resulting in the production of truncated proteins (Fig. 9A, E). The phenotypes of inbred B73 lines and the two maize mutants were analyzed after 4 days of growth under salt stress. Under normal conditions, the growth status of the wild type and mutants was not significantly different (Fig. 9). However, under 100 mM NaCl treatment, the shoot length of the *zmkea2* mutant was significantly lower than that of the wild type (*P*<0.05). The shoot dry weight of *zmkea2* was reduced greater than that of the wild type (Fig. 9B–D). Similarly, under 100 mM KCl condition, the shoot length of the *zmnhx5* mutant were significantly lower than that of the wild type (*P*<0.05). The shoot dry weight of *zmnhx5* was reduced greater than that of the wild type. ((Fig. 9F–H)). These results further verified that the *ZmNHX5* and *ZmKEA2* are important salt tolerance-related genes.

**Fig. 9.**
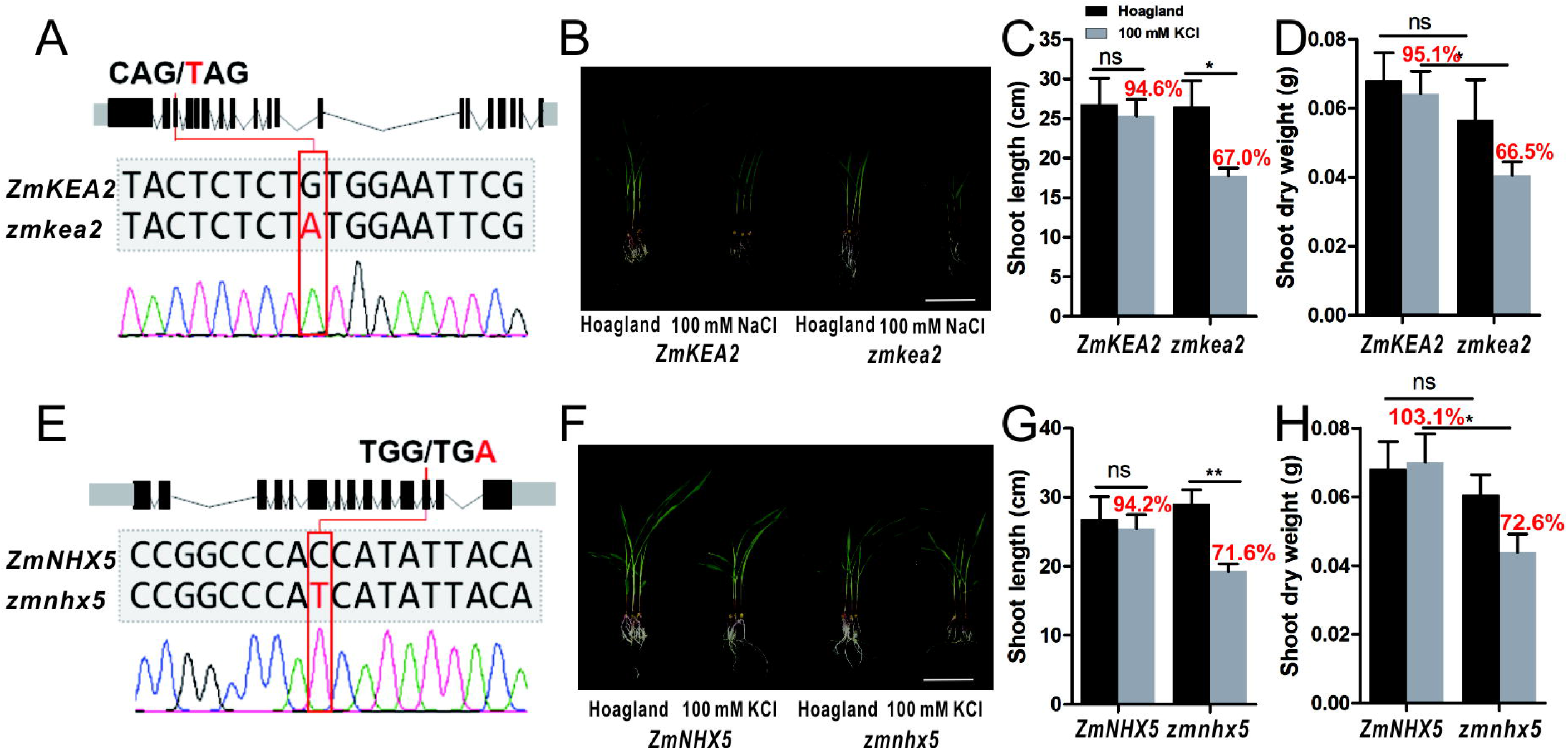
Growth performance of zmnhx5 and zmkea2 mutant seedlings under salt stress. Point mutation sites of EMS mutants zmnhx5 and zmkea2 were identified by PCR sequencing. Stop codons induced by EMS resulted in truncated proteins (a, e). Growth status (b, f), seedling length (c, g) and dry weight (d, h) of mutant seedlings shoot under 100 mM NaCl or under 100 mM KCl treatment for 4 days (n = 6, mean ± SEM). *, Significant difference (Student’s t-test; P < 0.05). NS, non significant.

## 3. Discussion

The cation/proton antiporter superfamily is responsible for monovalent cations exchange in fungi, bacteria, animals, and plants [12, 16]. Functional analyses of *CPA* genes have been conducted in *Arabidopsis* [22, 23], rice [17], wheat [24] and *Arachis hypogea* [25]. Most of them acted on maintaining the pH, Na^+^ and K^+^ homeostasis in plants under high salinity [26]. In maize, the *ZmNHX7* is homologous to *AtNHX7/SOS1*, which is critical for excluding Na^+^ from plant cell. The expression of *ZmNHX7* was up-regulated upon NaCl stress, and it is associated with the major quantitative trait locus (QTL), *qST1*, which confers salt tolerance on maize plants [27, 28]. Till now, there are few studies on maize *CPAs* gene family, and the functional analysis of *ZmCPAs* are rarely performed.

In this study, 35 *ZmCPA*s were identified in maize, and they were named according to previous reports [29] or based on their sequence similarity to known plant *CPA* genes. Analysis of chromosomal distribution revealed that *ZmCPA*s are evenly distributed on the 10 chromosomes of maize. Similarly, *CPA* genes are located on all chromosomes in wheat [24], and 15 out of 17 chromosomes in pear [30]. A phylogenetic tree using CPA proteins from *Arabidopsis*, rice and maize revealed that the homologous proteins from maize, *Arabidopsis* and rice were tightly clustered. For example, the *ZmNHX7, AtNHX7* and *OsNHX7* clustered together; the *ZmKEA3, AtKEA3* and *OsKEA3* gathered together; the *ZmCHX1, AtCHX1* and *OsCHX1* were in one clade. In *Arabidopsis*, genes with similar functions are clustered nearby, e.g., the *AtNHX1–4* localized on vacuolar membrane, *AtNHX5–6* for vesicle trafficking and *AtNHX7–8* localized on PM were distributed together, respectively. Thus it was speculated that the clustered homologous genes from maize and *Arabidopsis* or clustered maize genes had similar functions. Consistent with previous studies in *Arabidopsis* and pear, we also classified the CPA superfamily proteins into NHX, KEA and CHX subfamilies and into the N1–N3, K1–K2, and C1–C3 groups [12, 30].

The predicted sub-cellular localization of these proteins in different species is consistent to some extent. The *AtNHXs* exhibit vacuole, endosome, and PM localization [4] and the *ZmNHXs* have similar predicted localizations. Like *AtKEA1, AtKEA2* and *AtKEA3*, the *ZmKEA4* was predicted to localize to the chloroplast. All *ZmCHX* proteins were predicted to localize to the PM, like *AtCHX13* and *AtCHX14* [31]. The *ZmNHX5* and *ZmCHX17* were selected for transient expression in maize protoplast cells. As predicted, they were localized to the membrane system.

We analyzed the expression patterns of *ZmCPA* genes according to the NimbleGen maize microarray data. The *ZmNHX* and *ZmKEA* genes showed high expression at multiple developmental stages, suggesting that they are important for maize growth and development. The *ZmCHXs* were specifically expressed in anthers, indicating that they participate in reproductive organ development. Similar expression trends of *CPA* genes have been found in other plant species. For example, in wheat, *TaNHX2, TaNHX5* and *TaNHX8* are highly expressed at multiple developmental stages, and most *TaCHX* family genes are relatively highly expressed in anthers [24]. The expression patterns of selected genes in maize were verified by semi-qRT-PCR in this study.

The differential expression of *ZmCPAs* between the control and salinity treatments were analyzed using qRT-PCR. Under 100 mM NaCl treatment, the *ZmNHX5* and *ZmKEA2* were up-regulated in both leaf and root, providing evidence of their roles in NaCl stress in maize. Under 100 mM KCl treatment, the *ZmNHX2, ZmNHX5, ZmNHX6, ZmNHX8, ZmKEA1, ZmKEA2, ZmKEA5* and *ZmKEA6* were up-regulated in the leave. The up-regulated expression of them to K^+^ stress indicates that they might be involved in maintaining K^+^ homeostasis in maize. The *ZmNHX5, ZmKEA2, ZmCHX2, ZmCHX3, ZmCHX5* and *ZmCHX17* and were respectively introduced into the yeast mutant strain AXT3K. All constructs restored Na^+^-resistance of the AXT3K mutant. Furthermore, the growth reduction of maize EMS mutants under salt stress validated the functions of *zmnhx5* and *zmkea2* in salt tolerance. These analyses provided evidence for the role of CHX and KEA subfamilies in salt tolerance in maize. Similarly, *AtNHX5* and *AtNHX6* recovered salt tolerance in yeast [23], indicating that *ZmNHX*s have a common mode of action as other plant species. Unlike *ZmCHX* and *ZmKEA* from maize, neither *AtKEAs* nor *AtCHXs* from *Arabidopsis* are shown to improve yeast growth under high salt condition [4, 13].

It is worth mentioning that both *ZmNHX5* and *ZmKEA2* played an important role in salt tolerance of maize according to the results obtained in this study. The expression of *ZmNHX5* and *ZmKEA2* were up-regulated under salt stress (NaCl and KCl), and they can restore the Na^+^-resistance of yeast mutant. Moreover, the salt tolerance function (NaCl) of *ZmNHX5* and *ZmKEA2* were further verified by maize mutant experiment. The functional analysis of these two genes needs further study, and they are likely to be new gene targets for breeding salt tolerance maize.

## 4. Conclusion

We identified and characterized the *ZmCPA* superfamily, consisting of *ZmNHX, ZmKEA* and *ZmCHX* subfamily genes, from the maize genome. Gene and protein sequences analyses demonstrated high conservation of evolutionarily related members in each subfamily, while members differed among different groups. The existence of many helices, coils and TM regions in protein structure are consistent with the hydrophobic and membrane-bound nature of these proteins. Expression analyses revealed diverse expression patterns in various tissues and under different abiotic stresses, indicating that these genes are important for maize growth, development, and salt stress responses. Characterization of *ZmNHX5, ZmCHX2*, -*3*, -*5*, -*17* and *ZmKEA2* in yeast confirmed their roles in maintaining monovalent cation homeostasis and in salt stress tolerance. The functions of *ZmNHX5* and *ZmKEA2* were verified by phenotypic analyses of EMS mutants. This study not only describes numerous features of *ZmCPA* genes, but also provides the foundation for functional validation of these genes in further studies.

## 5. Materials and methods

### 5.1. Identification and bioinformatic analysis of ZmCPA gene superfamily

The known *CPA* genes in *Arabidopsis* were used to query the maize AGPv4 gene set (https://download.maizegdb.org/Zm-B73-REFERENCE-GRAMENE-4.0/) using a local BLASTP program with an E-value <1e-10. The putatively identified sequences were further confirmed by an HMMER search for the presence of signature Na^+^/H^+^ exchanger (PF00999) domain.

The phylogenetic tree was constructed using full-length CPA protein sequences of maize, *Arabidopsis*, and rice. Sequences were aligned using MUSCLE v3.8.31 [32, 33], and a phylogenetic tree was built using the neighbor-joining (NJ) method with MEGA 6.0 [34] with a bootstrap value of 1000 and amino acid substitution according to the Poisson model. The TM domains were predicted using tools at the TMHMM Server v. 2.0 (http://www.cbs.dtu.dk/services/TMHMM/). Predotar [35], TargetP [36], and WoLF PSORT (https://www.genscript.com/wolf-psort.html), three *in silico* programs, were used to predict the organellar localization of ZmCPA proteins. Putative conserved motifs in maize CPA proteins were identified using the MEME Suite 5.1.1 (http://meme-suite.org/tools/meme) with the following parameters: motif length, 10–50 aa; maximum number of motifs, 15.

### 5.2. Gene structure analysis

The DNA and transcript sequences of *ZmCPA* genes obtained from the maize sequence annotation database MaizeGDB were used to design gene-specific PCR primers with Primer3 (http://primer3.ut.ee/). The DNA and cDNA sequences were validated using PCR and RT-PCR with B73 genomic DNA and total RNA as templates. The gene-specific primers shown in Table S2. Alignment of validated DNA and cDNA sequences of each maize *CPA* gene revealed the structure of *ZmCPA* genes. Gene structure display server (GSDS 2.0) was used to display the exon–intron structure, and intron phases [37].

### 5.3. Plant materials and treatments

The maize B73 inbred line was used in this study. For qRT-PCR, the sterilized seeds were grown in a hydroponic system as described previously [38] with sterile water in a greenhouse at 27/23°C under a 12-h light/12-h dark photoperiod. Four days later, the plants were incubated in 1× Hoagland’s solution (PhytoTechnologies Ltd., Shawnee Mission, KS, USA) until the first leaves were fully expanded. Maize plants were grown with 1 × Hoagland’s solution as the control (CK) and with 1 × Hoagland’s solution containing 100 mM NaCl for the NaCl stress treatment or 100 mM KCl for the KCl stress treatment. These concentrations were maintained until the end of the experiments.

For maize EMS mutants experiment, the sterilized seeds were cultured hydroponically in a greenhouse at 27/23°C under a 12-h light/12-h dark photoperiod as described above. The 1× Hoagland’s solution was exchanged every 2 days. Ten days later, 100 mM KCl and 100 mM NaCl were added. After 4 days of salt stress, phenotypic analyses were performed.

### 5.4. Expression analysis of ZmCPA genes in different tissues

To investigate the spatiotemporal expression patterns of *ZmCPA* genes, the log2-transformed and RMA-normalized data for *ZmCPA* genes were downloaded from PLEXdb (http://www.plexdb.org/) [39]. A heat map was produced using the Lianchuan Bio Cloud Platform (https://www.lc-bio.cn/overview).

### 5.5 RNA isolation and cDNA synthesis

Total RNA was isolated from different tissues of the B73 inbred line, including seedling roots, leaves, 5-cm ears, immature tassels, anthers, pollens, silks, and seeds at 20 days after pollination, using Trizol reagent (Invitrogen, Carlsbad, CA, USA) according to the manufacturer’s protocol. All RNA was purified using DNase I (Thermo Scientific, Waltham, MA, USA). First-strand cDNA was synthesized from 1 μg total RNA (20 μL reaction volume) using a PrimeScript™ 1st Strand cDNA Synthesis Kit (Takara, Dalian, China) according to the manufacturer’s protocol.

### 5.6. Semi-quantitative reverse transcription PCR

All gene-specific primers were designed as shown in Table S5. Specific primers were used to amplify maize *Actin1* (*GRMZM2G126010*) as an internal control. Reactions were performed with 2 × Taq Master Mix (Vazyme, Nanjing, China) on a Bio-Rad Thermal Cycler (Bio-Rad, Hercules, CA, USA) using the following procedure: 5 min at 94 °C to start; 33 cycles of 30 s at 94 °C, 30 s at 59 °C and 2 min at 72 °C; and a final extension step of 72 °C for 10 min to complete the reaction. *Actin1* was amplified with 29 PCR cycles. Each PCR was performed in triplicate, mixtures without a template were employed as negative controls, and the maize *Actin1* amplicon served as an internal control for each gene investigated.

### 5.7. Real-time PCR

Real-time PCR was performed using TB Green Premix Ex Taq™ II (Takara, Dalian, China). An ABI 7500 real-time PCR system (Applied Biosystems, Foster City, CA, USA) was used with the following thermal cycling conditions: 95 °C for 5 min, followed by 40 cycles of 95 °C for 15 s, 60 °C for 5 s, and 72 °C for 34 s. The maize *Actin1* gene (*GRMZM2G126010*) was used as an endogenous control to normalize the samples. Based on the cDNA sequences of *ZmCPA* genes, real-time PCR primers (Table S6) were designed with Primer 3 (http://primer3.ut.ee/). The experiment was performed with three technical replicates for each sample. The specificity of each PCR reaction was confirmed by melting curve analysis of the amplicons. The comparative 2^− ΔΔ^CT method was used to calculate the relative transcript levels in the samples [40].

### 5.8. Subcellular localization

The full-length cDNAs of *ZmCHX17* and *ZmNHX5* were amplified using the primers listed in Table S7 and then were introduced into the pM999-EGFP vector to construct the GFP fusion proteins ZmCHX17-GFP and ZmNHX5-GFP, respectively, with the ClonExpress II One Step Cloning Kit (Vazyme). Constitutive expression of the fused constructs, ZmCHX17-GFP and ZmNHX5-GFP, was driven by the cauliflower mosaic virus 35S promoter. Maize mesophyll protoplasts were isolated and prepared from etiolated leaves according to established protocols [41]. The plasmids harboring the ZmCHX17-GFP and ZmNHX5-GFP fusion constructs were co-transfected with the OsSCAMP1-RFP construct into protoplast cells, respectively. OsSCAMP1 is a known rice secretory carrier membrane protein and was used as a membrane protein control [18]. The transformed protoplast cells were cultured at room temperature overnight and then observed under a Leica SP8 confocal microscope (Leica, Wetzlar, Germany).

### 5.9. Functional expression of ZmCPAs in yeast

The coding sequences of *ZmNHX5, ZmCHX2, ZmCHX3, ZmCHX5, ZmCHX17*, and *ZmKEA2* were each cloned into the PDR196 vector, and then transformed into the yeast strain AXT3K (*ena1-4::HIS3,nha1::LEU2, nhx1::KanMX*). The transformed yeast cells were cultured overnight at 29 °C in YPDA medium containing 1 mM KCl. Cell cultures were diluted in water to an A_600_ of 0.8. To test cation tolerance, 5-μL aliquots from yeast cultures or 10-fold serial dilutions were spotted onto AP [42] plates containing 1 mM KCl and supplemented with or without 20 mM NaCl.

## List of abbreviations

CPA: monovalent cation proton antiporter;
NHX: Na^+^/H^+^ exchanger;
KEA: K^+^ efflux antiporter;
CHX: cation/H^+^ exchanger;
Na^+^: sodium;
Cl^-^: chloride;
TM: transmembrane;
K^+^: potassium;
H^+^: proton;
MW: molecular weight;
pI: isoelectric point;
RT-PCR: reverse transcription polymerase chain reaction;
DNA: deoxyribonucleic acid;
cDNA: complementary DNA;
NJ: neighbor-joining;
aa: amino acid;
kDa: kilodalton;
TF: transcription factor;
semi qRT-PCR: semi-quantitative reverse transcription polymerase chain reaction;
AP: Arg phosphate;
EMS: ethyl methanesulfonate;
MEMD: Maize EMS-induced Mutant Database;
QTL: quantitative trait locus;
GFP: green fluorescent protein;
RFP: red fluorescent protein.

## Authors’ contributions

Y.T., Y.Z., (Yanxin Zhao) and J.Z. conceived the experiment. M.K, M.L, J.L, Y.Z. (Yunxia Zhang), Z.F, W.S, R.Z, R.W., and Y.W. performed bioinformatic analysis and data acquisition. M.K, M.L, and Y.Z (Yanxin Zhao) analyzed the data and wrote the manuscript. All authors revised and approved the final manuscript, and agreed to be accountable for this work.

## Acknowledgements

We thank Prof. Liwen Jiang (Chinese University of Hong Kong) for kindly providing pM999-EGFP and membrane protein marker (OsSCAMP1-RFP) plasmids, Prof. Xiaoduo Lu (Qilu Normal University) for identifying and providing maize EMS mutants *zmkea2* and *zmnhx5*, and Profs. Yi Wang and Zhenxian Zhang (China Agricultural University) for kindly sharing yeast mutant AXT3K and yeast expression vector pDR196.

